# Bi-directional Impacts of Heterotypic Interactions in Engineered 3D Human Cardiac Microtissues Revealed by Single-Cell RNA-Sequencing and Functional Analysis

**DOI:** 10.1101/2020.07.06.190504

**Authors:** Tracy A. Hookway, Oriane B. Matthys, David A. Joy, Jessica E. Sepulveda, Reuben Thomas, Todd C. McDevitt

## Abstract

Technological advancements have enabled the design of increasingly complex engineered tissue constructs, which better mimic native tissue cellularity. Therefore, dissecting the bi-directional interactions between distinct cell types in 3D is necessary to understand how heterotypic interactions at the single-cell level impact tissue-level properties. We systematically interrogated the interactions between cardiomyocytes (CMs) and cardiac non-myocytes in 3D self-assembled tissue constructs in an effort to determine the phenotypic and functional contributions of cardiac fibroblasts (CFs) and endothelial cells (ECs) to cardiac tissue properties. One week after tissue formation, cardiac microtissues containing CFs exhibited improved calcium handling function compared to microtissues comprised of CMs alone or CMs mixed with ECs, and CMs cultured with CFs exhibited distinct transcriptional profiles, with increased expression of cytoskeletal and ECM-associated genes. However, one month after tissue formation, functional and phenotypic differences between heterotypic tissues were mitigated, indicating diminishing impacts of non-myocytes on CM phenotype and function over time. The combination of single-cell RNA-sequencing and calcium imaging enabled the determination of reciprocal transcriptomic changes accompanying tissue-level functional properties in engineered heterotypic cardiac microtissues.

## Introduction

Engineered cardiac tissue models have moved towards incorporating multiple cell populations in order to more accurately mimic the cellular complexity of the native heart. Contractile cardiomyocytes (CMs), the primary (i.e. “parenchymal”) cells of the heart, are supported by various non-parenchymal cell types, most notably endothelial cells (ECs) and cardiac fibroblasts (CFs). The significance of non-parenchymal cells to cardiac function is reflected by the fact that although CMs constitute approximately 75% of the mature heart by volume, they make up less than one-third of the total number of cells in the heart^1–4^.

Endothelial cells, originating primarily from lateral plate mesoderm, are the most abundant non-myocyte population in the adult heart^5–7^. ECs communicate with CMs primarily via paracrine and endocrine signaling (i.e. release of platelet-derived growth factor-B, neuregulin, and nitric oxide), and support CM metabolism, survival, and contractility^8,9^. Despite the high tissue density and close proximity of ECs to CMs, ECs are physically separated from CMs by a thin basement membrane^10^, explaining why ECs primarily influence CM fate and function via secreted molecules, and not as a result of direct cell-cell interactions.

Cardiac fibroblasts, on the other hand, are in intimate physical contact with multiple neighboring CMs and constitute ~15% of non-myocytes in the heart^5^. CFs arise during embryonic development from epicardial cells and endocardial second heart field progenitors that undergo an epithelial-to-mesenchymal transition, as well as from a small population generated in the neural crest that primarily populates the outflow tract of the heart^11–13^. CFs derived from the epicardium constitute the majority of the fibroblast population (80–85%) in the heart^12,13^, and are typically found in the ventricular myocardium, resulting in a close spatial relationship between CFs and CMs, such that every CF is surrounded by multiple CMs^14,15^. As a result of their physical juxtaposition, CM-CF communication is mediated largely by direct cell-cell interactions, such as physical intercellular adhesions and gap junctions^16^. CFs also contribute to cardiac tissue structure, function, and homeostasis through the secretion of growth factors and signaling molecules (i.e. fibroblast growth factor 2, transforming growth factor-B, interleukin-1B, interleukin-6)^17^, as well as extracellular matrix (ECM) molecules and proteases responsible for cardiac remodeling^18^. Ontogenic differences in CF morphology and ECM have also been determined between isolated fibroblasts from fetal and adult cardiac tissue. Fetal CFs tend to be smaller, more proliferative, and produce ECM rich in fibronectin, periostin, and heparin-binding EGF-like growth factor, which supports CM proliferation^18–20^. In contrast, adult CFs are larger, less proliferative, and produce more structural ECM (i.e. collagen and elastin) to support CM hypertrophy^19,20^.

The ability to efficiently generate CMs from human pluripotent stem cells (hPSCs) has enabled broad use of these cells as a source for tissue engineering efforts. The majority of cardiac tissue engineering strategies acknowledge the need for including multiple heterogeneous cardiac cell populations in order to better model the cellular composition and physiology of the native tissue. However, although inclusion of a nonmyocyte population is necessary for robust tissue formation and long-term maintenance of engineered cardiac tissues^21^, the specific mechanisms by which non-myocytes impact CMs have yet to be elucidated. This lack of clarity is confounded by the fact that the wide variety of non-myocytes that have been paired with hPSC-derived CMs are largely undefined populations derived from different sources (i.e. primary human-derived vs. stem cell-derived) of varying ages^21^. Further heterogeneity arises from the hPSC-CM differentiations, which yield contaminating fractions of non-target cell populations as well as cells of varying levels of commitment/maturity.

Single-cell RNA-sequencing has emerged as a powerful tool to dissect multicellular heterogeneity throughout tissue development^22,23^ or within heterogeneous hPSC-differentiations^24,25^. Applying single-cell RNA-seq to engineered heterotypic tissue constructs enables the determination of multicellular phenotypes at single-cell resolution within the context of tissue-level properties. Therefore, in this study we employed a systematic approach to interrogate the specific contributions of cardiac-specific nonmyocytes to cardiomyocyte phenotype and cardiac tissue function, by determining bidirectional single-cell transcriptomic changes that accompany functional differences between engineered cardiac tissue constructs.

## Results

### Non-myocytes enable rapid microtissue compaction

To explore the effects of non-myocytes on cardiac microtissue formation and function, we compared homotypic microtissues formed from CMs alone to heterotypic cardiac microtissues formed from CMs+endothelial cells (CM+EC), CMs+fetal cardiac fibroblasts (CM+fCF), and CMs+adult cardiac fibroblasts (CM+aCF) (Figure 1A). These cardiac cells self-assembled^26^ into 3D microtissues within 24h after seeding into inverted pyramidal microwells (Figure 1Bi); however, all heterotypic microtissues compacted to form spheroids more rapidly than homotypic tissues (Figure 1Bii,C). After 7 days of culture, CM alone microtissues were of similar size to heterotypic microtissues (Figure 1Biii) and all groups continued to decrease in tissue diameter (Figure 1C), but the heterotypic microtissue groups displayed a dark core at the center that was absent in the CM alone microtissues (Figure 1Biii). After 30 days of culture, all microtissue groups had decreased in size (Figure 1C), but there were no major differences in microtissue size or morphology between conditions (Figure 1Biv).

**Figure 1.**
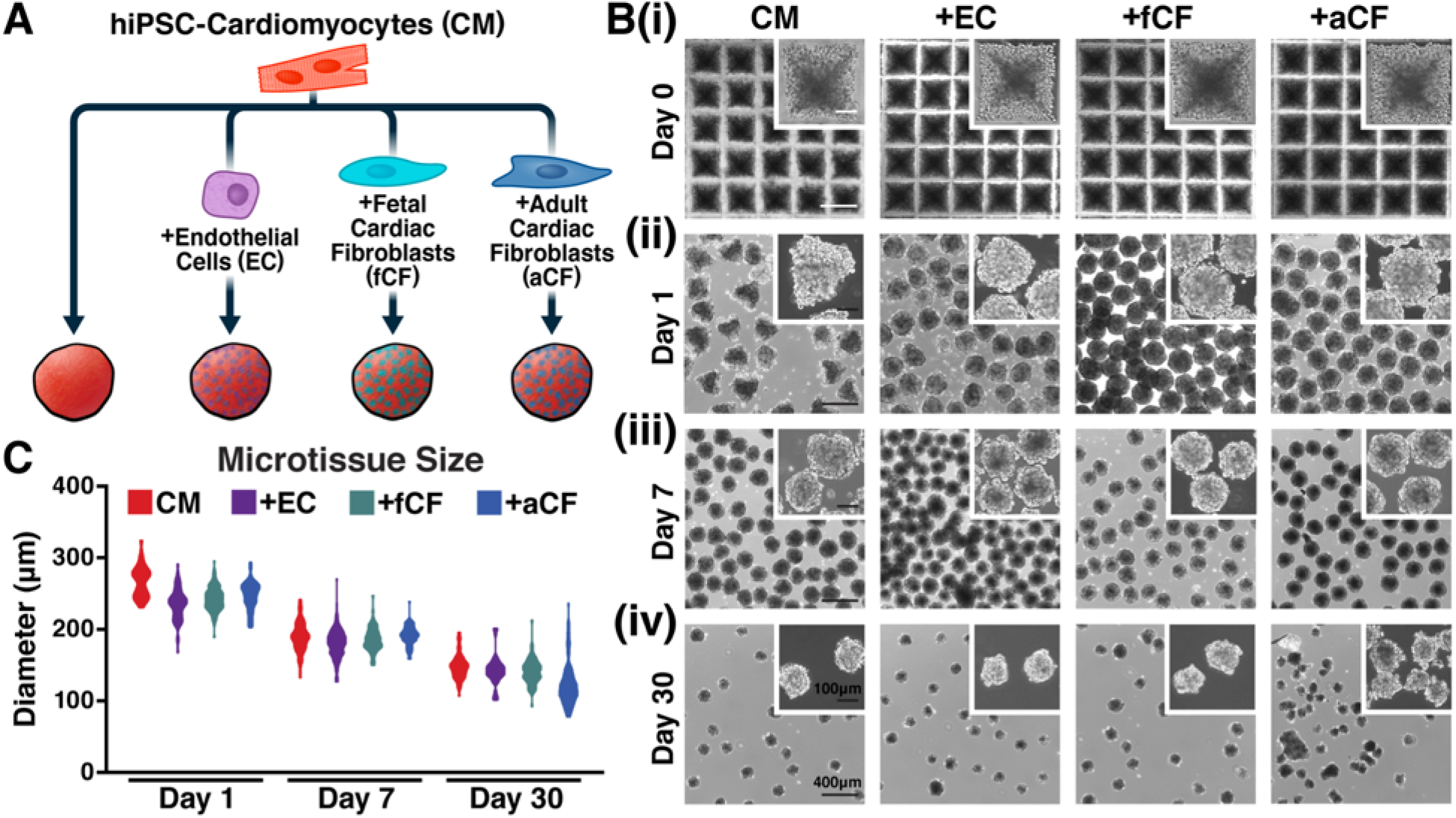
Heterotypic cardiac microtissue formation. (A) Schematic of cardiac microtissue conditions. (B) Phase contrast imaging of cardiac microtissues throughout formation and culture duration. Microtissues self-assembled within 24h and compacted over the course of 30 days of culture. Heterotypic microtissues compacted more rapidly than homotypic CM alone tissues. Scale bar = 400μm (inset = 100μm). (C) Diameter of all cardiac microtissue groups decreased throughout culture duration.

### Non-myocytes form a core within heterotypic cardiac microtissues

Hematoxylin and eosin staining of the microtissues revealed relatively uniform, high cell densities, regardless of heterotypic pairings (Figure 2Ai, 2Bi). Therefore, to characterize the multicellular organization of heterotypic cell populations, we stained the microtissues for specific phenotypic markers. At day 7, most of the heterotypic microtissues contained a core of non-myocytes, based upon the absence of cardiac troponin T (cTnT) expression in regions that were positive for wheat germ agglutinin (WGA; Figure 2Aii), which correlated to the dark centers observed in the phase contrast images (Figure 1Biii). After 30 days of culture, non-myocyte cores were only evident in a subset of the heterotypic microtissues, primarily those containing fetal CFs (Figure 2Bii). To determine whether culture with non-myocytes impacted the phenotypic maturation of the CMs, we stained for various isoforms of troponin I. Seven days post tissue formation, expression of cardiac troponin I (cTnI) expression—a marker of CM maturity compared to slow/fast skeletal troponin I (sTnI)—was more abundant in microtissues containing CFs and localized adjacent to the non-myocyte cores (Figure 2Aiii). However, after 30 days, there were less apparent differences in cTnI expression between experimental groups (Figure 2Biii).

**Figure 2.**
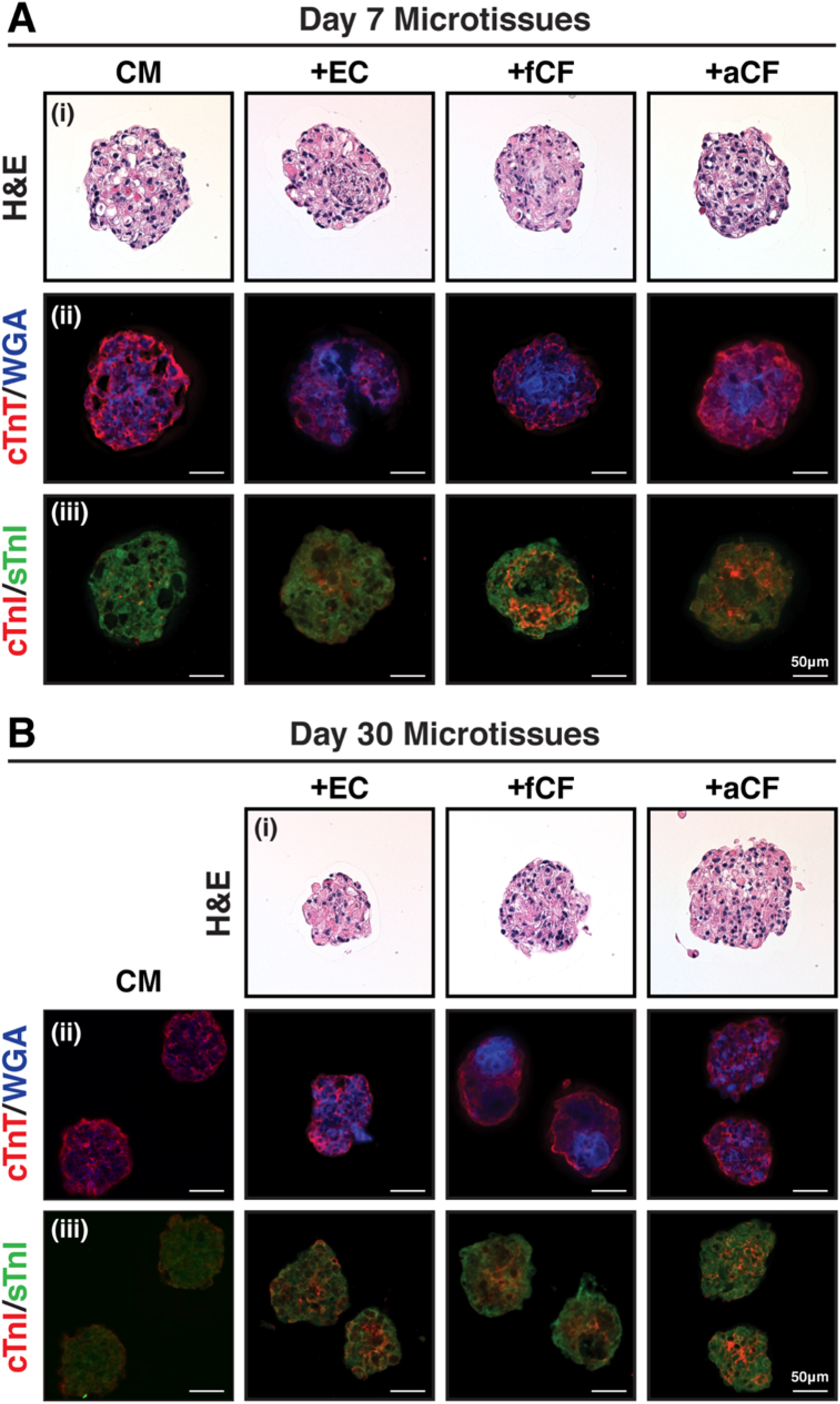
Histology and immunostaining of microtissue sections. Day 7 (A) and day 30 (B) cardiac microtissue sections stained with hematoxylin and eosin (i), cardiac troponin T (cTnT) and wheat germ agglutinin (WGA) (ii), and cardiac troponin I (cTnI) and slow+fast troponin I (sTnI) (iii). Day 7 heterotypic microtissues contained cores of non-myocytes (WGA+/cTnT-) that were bordered by higher cTnI expression. Scale bar = 50μm.

### Cardiac fibroblasts accelerate calcium handling properties of cardiac microtissues

In order to assess the effect of heterotypic culture on microtissue function, we performed calcium imaging. Individual microtissues in each condition exhibited independent spontaneous beating, indicating that they functioned as autonomous tissue constructs (Figure 3A). Seven days after tissue formation, cardiac microtissues containing CFs (both fetal and adult) exhibited improved calcium handling profiles, characterized by increased amplitude (F/F_0_), and higher maximum upstroke and downstroke velocities than homotypic or CM+EC microtissues (Figure 3B). Microtissues cultured for 30 days exhibited higher amplitudes and faster stroke velocities than their day 7 counterparts (Figure 3C), demonstrating that culture duration improved calcium handling properties regardless of starting conditions. However, no distinction in transient profiles of the microtissues containing CFs compared to those with CMs alone or ECs was evident after 30 days, and instead, the CM alone microtissues exhibited calcium transients with the greatest amplitude and fastest stroke velocities.

**Figure 3.**
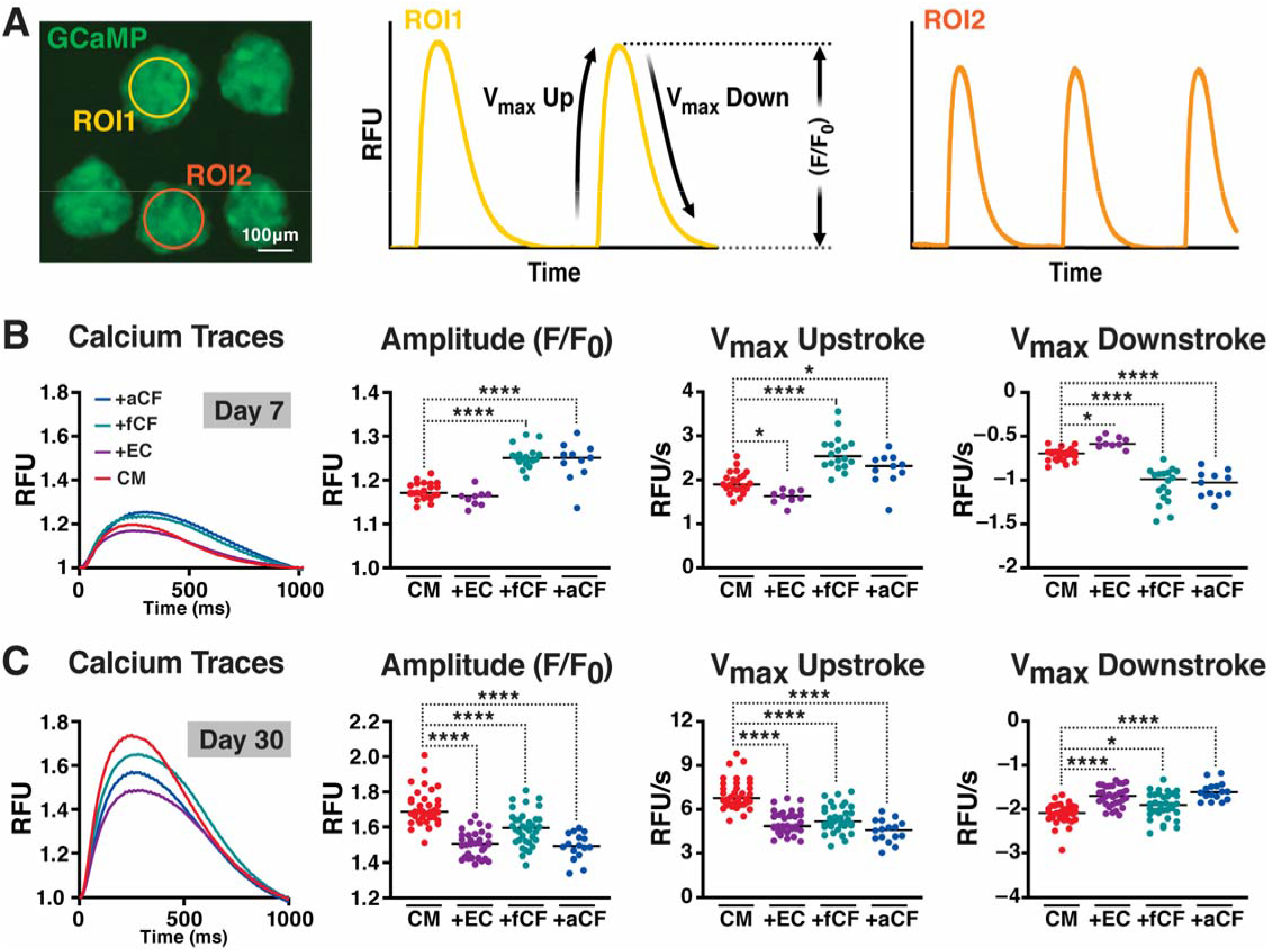
Cardiac microtissue calcium handling properties. (A) GCaMP fluorescence of cardiac microtissues and representative traces of spontaneous calcium transients with kinetic properties defined. Day 7 (B) and day 30 (C) cardiac microtissue calcium transient traces, amplitude values, and maximum upstroke and downstroke velocities (microtissues subjected to 1Hz stimulation). Day 7 microtissues containing fetal and adult CFs exhibited more mature calcium transients (higher amplitude and velocity values), but this distinction was attenuated at day 30. * p<0.05; ** p<0.01; **** p<0.0001.

To further assess calcium handling properties, we subjected day 7 and day 30 cardiac microtissues to a series of increasing electrical field stimulation frequencies, at 0.5Hz, 1Hz, 2Hz, and 4Hz (Figure 4Ai,Bi). At day 7, all microtissue groups were able to keep pace with 0.5Hz and 1Hz stimulation (Figure 4Aii). At 2Hz, all groups except CM+EC were able to keep pace with the stimulation, but all tissue groups failed to respond appropriately above 2Hz stimulation. Aligning the calcium transient traces to the input stimulation pulses revealed that the microtissues that failed to respond appropriately to the stimulation frequency instead responded at a lower, but harmonic beat rate (Supplementary Figure 1). At day 30, despite the improved calcium transient profiles (Figure 3C), the microtissues did not respond as robustly to the higher stimulation frequencies as their day 7 counterparts (Figure 4Bii). Most CM alone tissues (85.7%) and 100% of heterotypic microtissues were able to pace at 1Hz stimulation, yet only 20.6% of CM+fCF microtissues and 6.3% of CM+aCF tissues were able to respond to 2Hz stimulation. The lower harmonic response of these microtissues indicates that their calcium handling machinery could not take up and release calcium stores in time for higher pulse stimulation frequencies.

**Figure 4.**
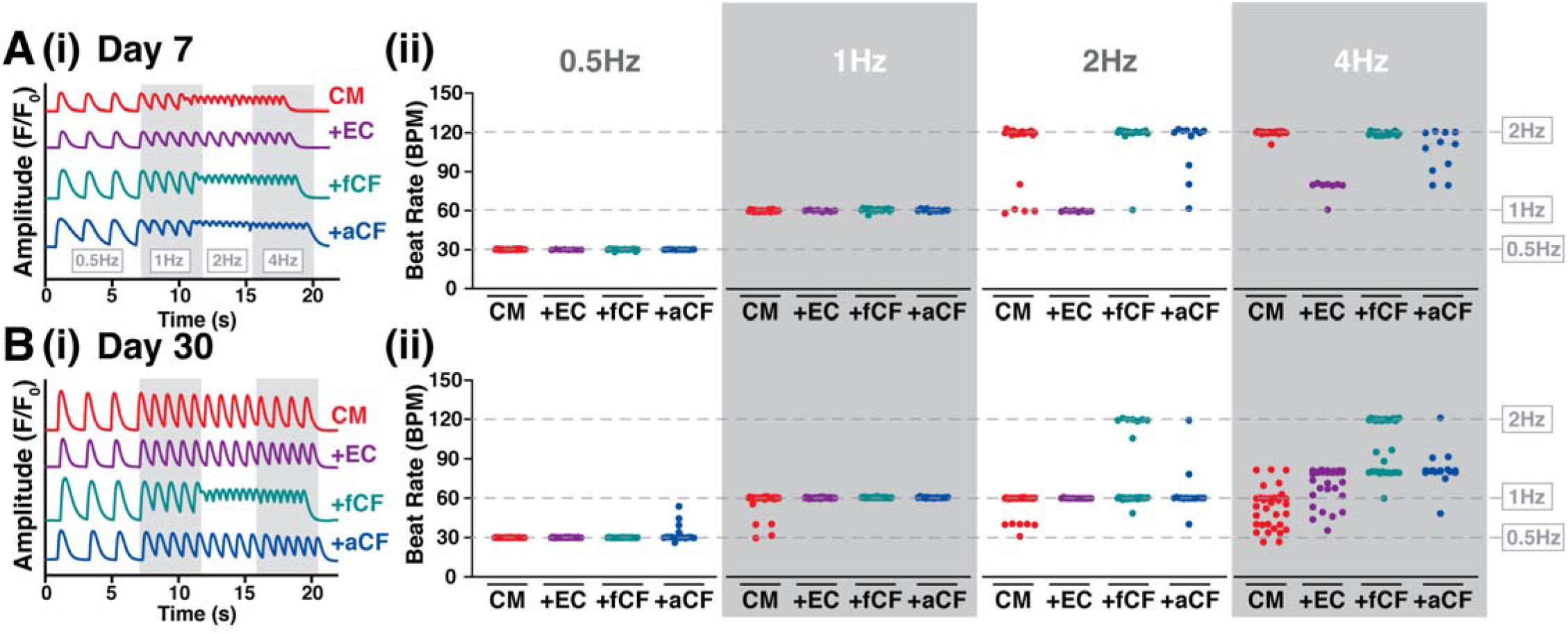
Calcium handling response to increasing electrical field stimulation. Day 7 (A) and day 30 (B) cardiac microtissue calcium transient traces (i) and responsive beat rate analysis (ii) at 0.5, 1, 2, and 4Hz stimulation frequencies. At day 7, microtissues containing ECs only respond to stimulation frequencies up to 1Hz, while the other microtissues groups are able to respond to 2Hz stimulation. At day 30, most tissues lost the ability to respond to 2Hz stimulation.

### Cardiomyocyte phenotype changes as a result of cardiac fibroblast co-culture

In order to determine the phenotypic influences of heterotypic interactions, we performed single-cell RNA-sequencing on cells from each culture condition and each time point (input day 0, day 7, and day 30; Figure 5A). Since flow cytometry methods are notoriously difficult to accurately sort cardiac cells due to lack of differential surface markers, we “sorted” cells using transcriptomic identification methods. A single cluster was identified as ECs based on expression of PECAM1, FLT, and KDR (4.7% cells total; Figure 5Bii,C) and two clusters were classified as CFs based on elevated expression of POSTN, FN1, and THY1 (8.8% cells total; Figure 5Biii,C). The remaining clusters were characterized as CMs based on expression of TNNT2, TNNI1, and ACTC1 (86.5% cells total; Figure 5Bi,C). A total of 18 clusters were assigned as CMs, suggesting that several different CM phenotypes existed in the pooled population of cells.

**Figure 5.**
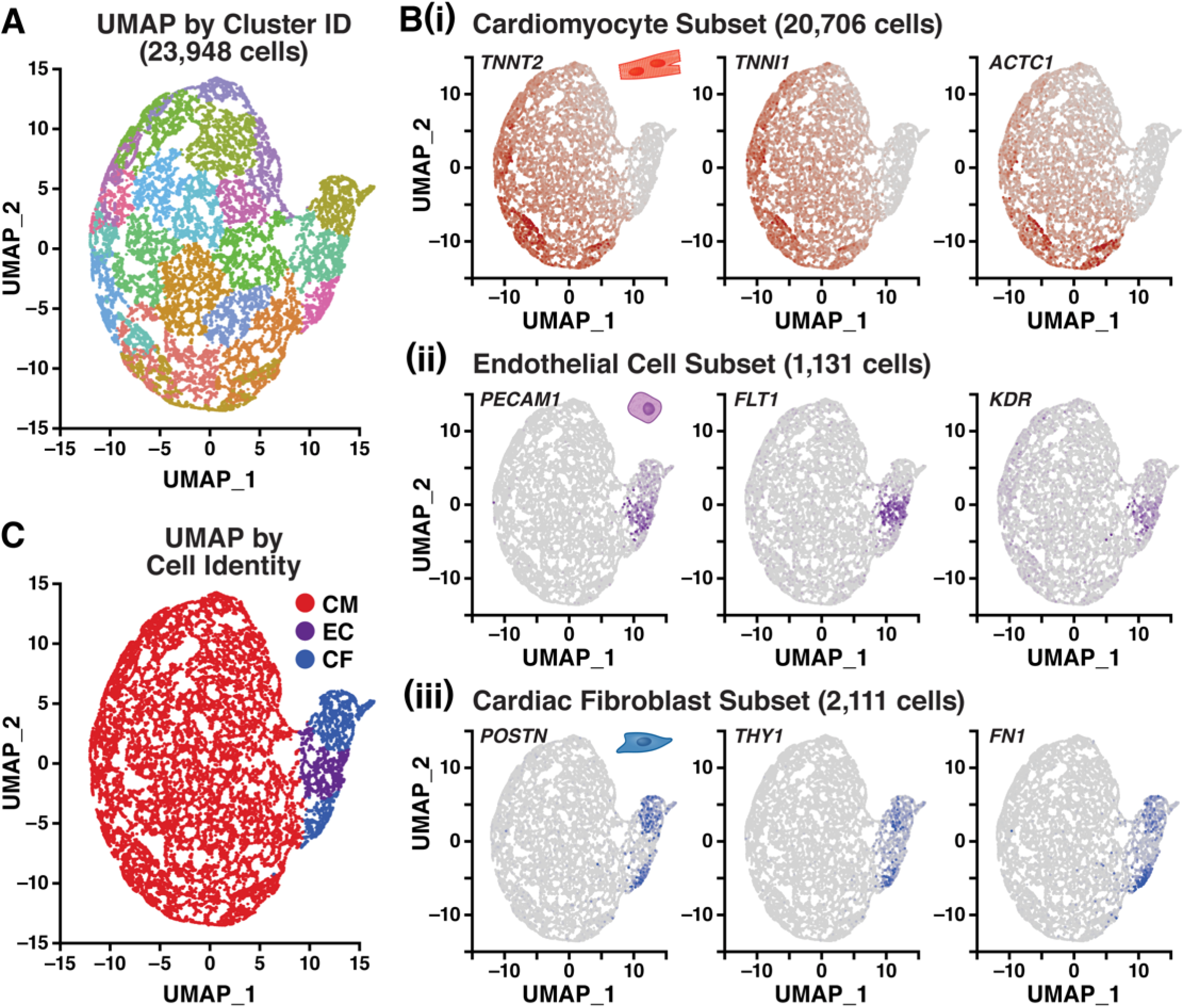
Single-cell RNA-sequencing analysis to identify cardiac cell types. (A) UMAP clustering of cells from all heterotypic pairings and culture time points. (B) Identification of cell types by expression analysis of TNNT2, TNNI1, ACTC1 for CMs (i), PECAM1, KDR, FLT1 for ECs (ii), and POSTN, THY1, FN1 for CFs (iii). (C) UMAP of all cells colored by cell type identity.

To interrogate these phenotypes, we performed secondary analysis by subsetting the cells classified as CMs, ECs, or CFs and separately re-clustering and analyzing differential gene expression within the different cell identities. Cells classified as CMs were subset and re-normalized using zinbwave reduction, resulting in 20 clusters that were separated largely along the axis of culture time (Figure 6A). Day 0 CMs represented ~15% of the analyzed CMs and clustered together regardless of initial heterotypic mixing, confirming a homogeneous input population. By day 7, however, CMs mixed with either fetal or adult CFs (4,182 CMs) were largely separated from CMs that were mixed with ECs or from homotypic cultures (4,233 CMs), highlighting a shift in CM phenotype as a result of CF co-culture for 7 days (Figure 6B). Yet, after 30 days of culture, CMs from all microtissue conditions (~44% of total CMs) clustered together regardless of heterotypic pairing, indicating a return to a more homogeneous CM phenotype after prolonged 3D culture.

**Figure 6.**
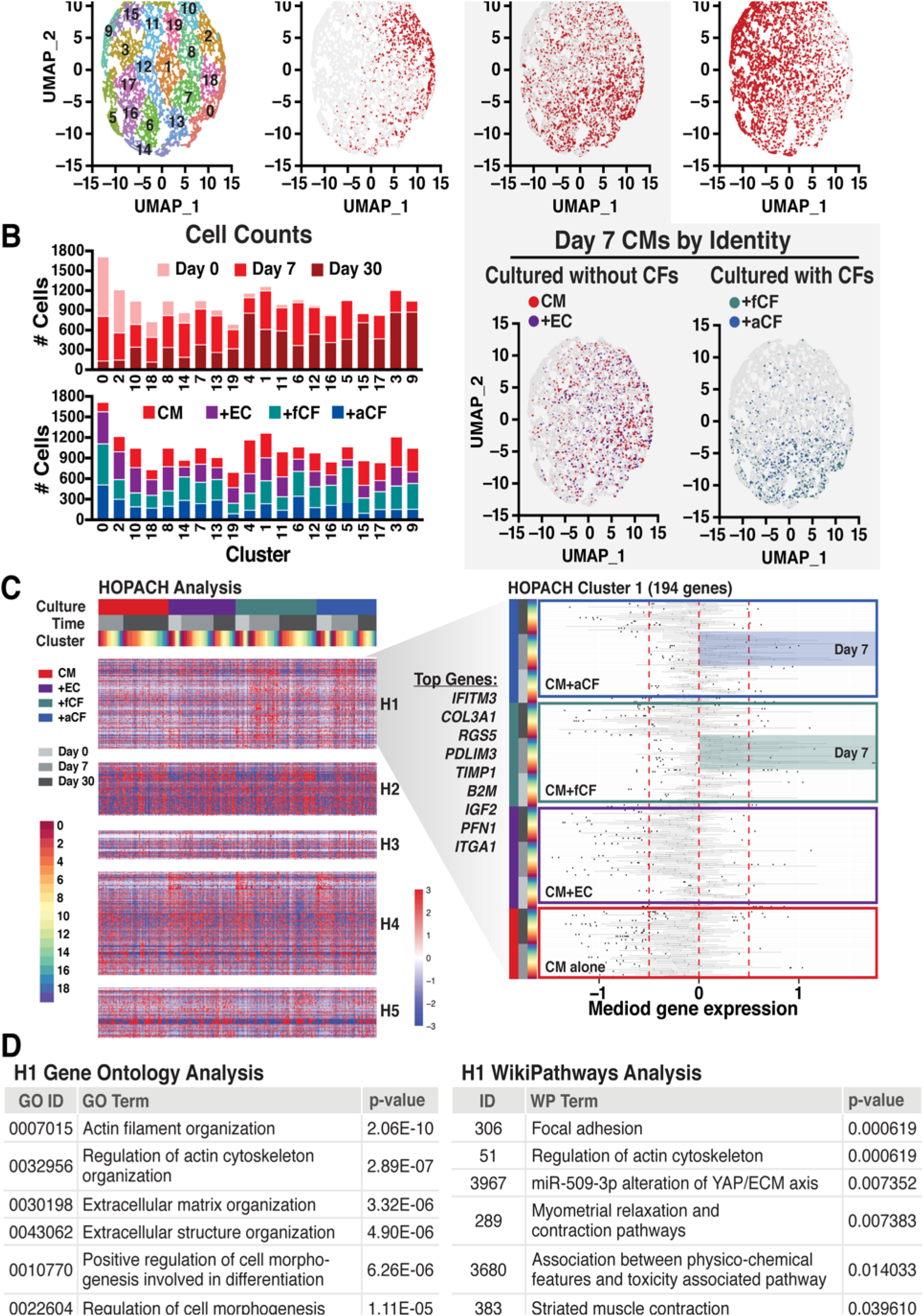
Cardiomyocyte subset analysis of single-cell RNA-sequencing data. (A) UMAP clustering of classified CMs colored by cluster identity and by culture time point. Day 7 CMs also colored by tissue condition to identify transcriptomic split between CMs cultured alone or with ECs and CMs cultured with fetal and adult CFs. (B) Cell counts for each UMAP cluster organized by time point and by tissue condition. (C) Heatmap of HOPACH clustering of CMs and plot of HOPACH cluster 1 (H1). (D) Gene ontology and WikiPathways results for H1.

We next employed Hierarchical Ordered Partitioning and Collapsing Hybrid (HOPACH) clustering to derive unbiased cell type-specific gene lists (H1_CM_-H5_CM_), followed by gene ontology (GO) and WikiPathways analysis. HOPACH identified the specific groups of CMs that contributed to the differential expression in each gene list, regardless of UMAP cluster, time point, or heterotypic pairing (Figure 6C, Supplementary Figure 2, Supplementary Table 1). For instance, H1_CM_ mirrored the UMAP clustering split of CMs that were cultured with CFs versus without. H1CM contained 194 genes which were upregulated in CMs from day 7 +fCF and +aCF cocultures, including *IFITM3, COL3A1, RGS5, PDLIM3, TIMP1, IGF2,* and *SPARC* (Figure 6C). These genes corresponded to ECM organization, cytoskeletal organization, and cell morphogenesis GO terms, as well as cytoskeletal, ECM, and physical contraction WikiPathways results (Figure 6D), indicating that the presence of CFs may lead to a CM phenotype that is more capable of remodeling its local environment. We confirmed upregulation of several of the top differentially-expressed genes between day 7 CMs cultured with CFs vs. without *(PDLIM3, RGS5, IGF2,* and *COL3A1)* by RNAscope (Figure 7).

**Figure 7.**
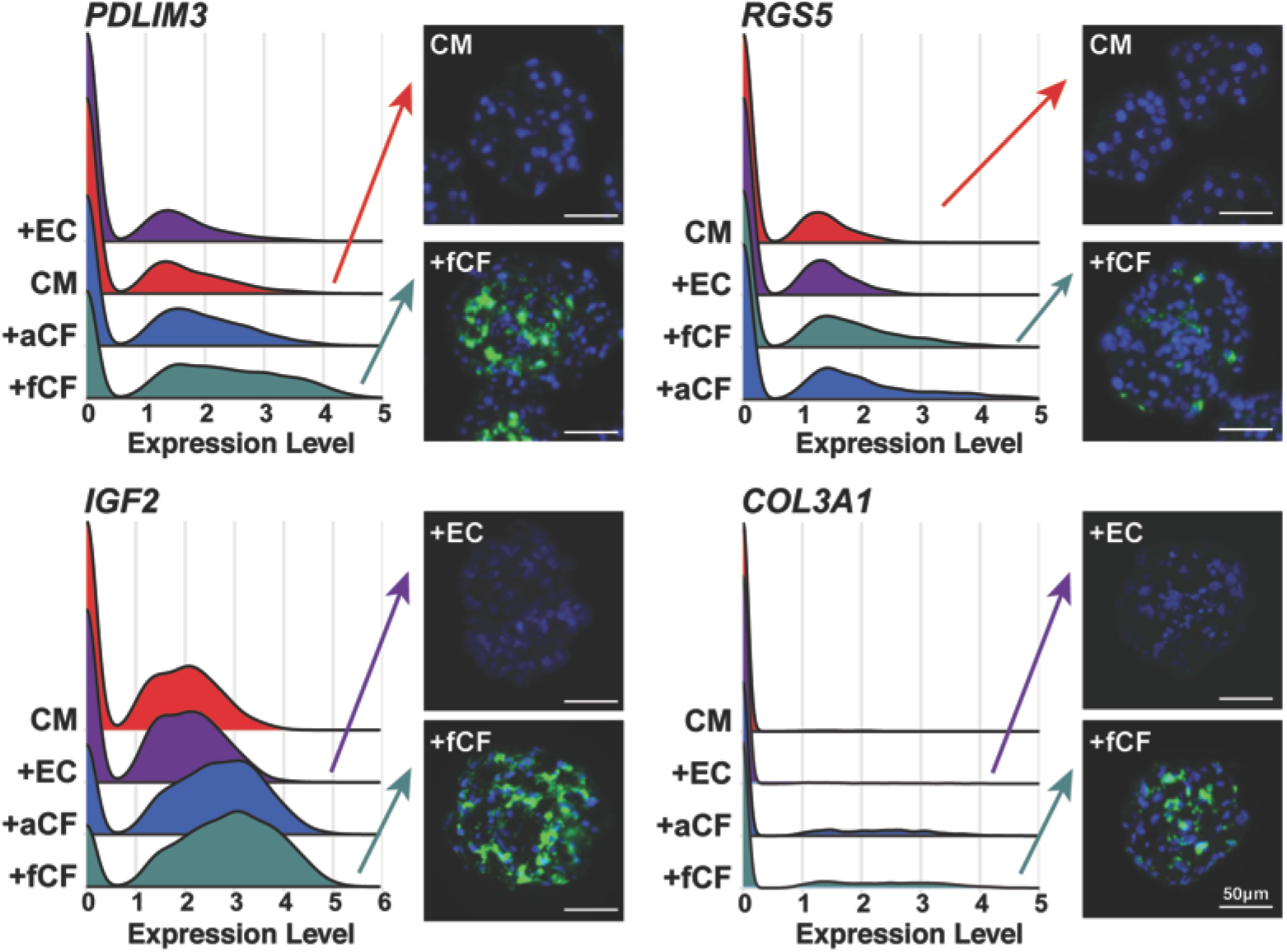
Validation of top differentially expressed genes in cardiomyocytes that were cultured with cardiac fibroblasts vs. without. In situ hybridization visualization with RNAscope validated higher mRNA expression of *PDLIM3, RGS5, IGF2,* and *COL3A1* in cardiac microtissues that contained CFs.

Other HOPACH clusters contained genes related to ribosomal processing (H2_CM_), cell cycle (H3_CM_), and mitochondria (H5_CM_) (Supplementary Figure 2). H5_CM_ contained genes with increased expression in day 30 CMs from UMAP clusters 4, 9, and 15 (increased expression in all microtissue groups), with GO terms that were largely relevant to cardiac function and development (Supplementary Figure 2), reflecting increased culture differentiation or maturation profiles.

### Non-myocyte phenotypic changes as a result of heterotypic co-culture

During initial cell identification after sequencing, 1,131 cells were identified as ECs based on a panel of markers (Figure 5B). However, only about 67% of these originated from an EC-containing microtissue group; the remainder came from CM+CF microtissues (Figure 8A,B). The majority of cells identified as ECs were from the initial (day 0) time point (~94%). However, HOPACH analysis and GO assessment identified several gene clusters (H1EC-H9EC) that changed expression patterns at day 7 and 30 as a result of heterotypic culture (Supplementary Figure 3, Supplementary Table 2). H1_EC_ contained 63 genes, such as *EDN1*, that were expressed at higher levels in input (day 0) ECs than in ECs post-3D-heterotypic culture (Figure 8C). H1_EC_ was enriched with genes annotated by GO terms related to neutrophil activation, muscle contraction, intracellular transport, and metabolic processes, as well as proteasomal- and glycolysis-related WikiPathways terms (Figure 8D). H3_EC_, on the other hand, contained genes that were upregulated in day 7 and day 30 ECs, including *KDR*, *IL32*, and *HYAL2* (Figure 8C). The relevant GO terms were related to endothelial development and migration as well as ECM organization and cell junction assembly, and WikiPathways results included cardiac calcium regulation and contraction pathways, indicating that heterotypic culture alters CM-EC interactions related to cardiac function (Figure 8D). The other HOPACH clusters listed differential gene expression in cells that did not come from the CM+EC microtissues, and therefore were not included in this secondary analysis (Supplementary Figure 3).

**Figure 8.**
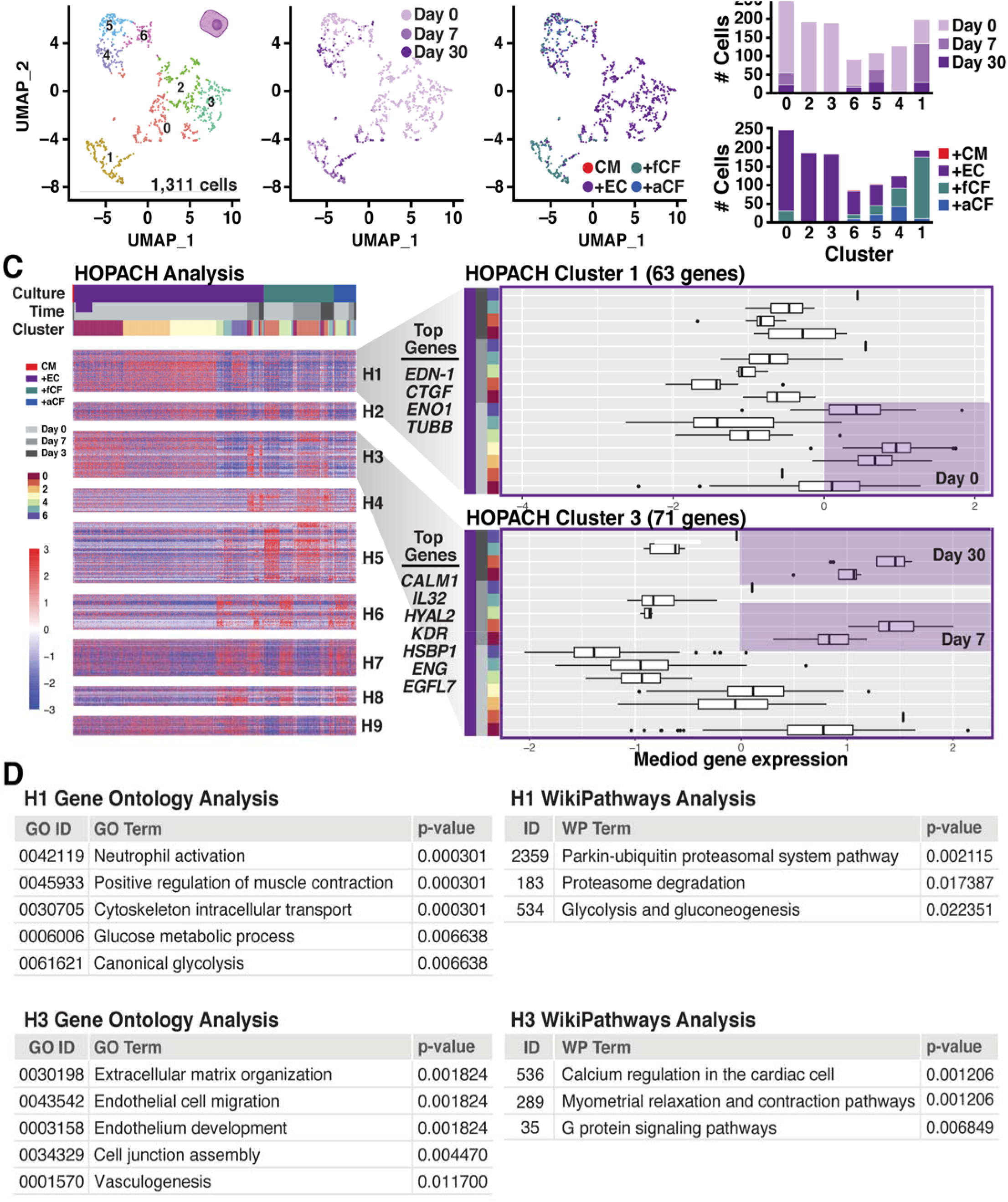
Endothelial cell subset analysis of single-cell RNA-sequencing data. (A) UMAP clustering of classified ECs colored by cluster identity, culture time point, and tissue condition. (B) Cell counts for each UMAP cluster organized by time point and by tissue condition. (C) Heatmap of HOPACH clustering of ECs and plot of HOPACH clusters 1 and 3 (H1 and H3, respectively). (D) Gene ontology and WikiPathways results for H1 and H3.

A total of 2,111 CFs were identified across all culture conditions and time points (Figure 5B), with only a small fraction of CFs (~6%) coming from the CM+EC and CM alone tissue groups, likely attributable to the stromal/fibroblastic cells derived from hiPSC-CM and hiPSC-EC differentiations. Of the identified CFs that came from the +CF microtissue groups, 73% were a result of the input cells (aCF and fCF at day 0), ~24% came from day 7 tissues, and only ~3% were identified after the full 30 days of culture (Figure 9A,B). Since immunostaining demonstrated the presence of non-myocytes in day 7 and 30 CM+CF microtissues (Figure 2A), the sharp decline in the numbers of CFs identified after 3D heterotypic co-culture is likely due to loss of the cells during microtissue dissociation rather than cell death or differentiation.

**Figure 9.**
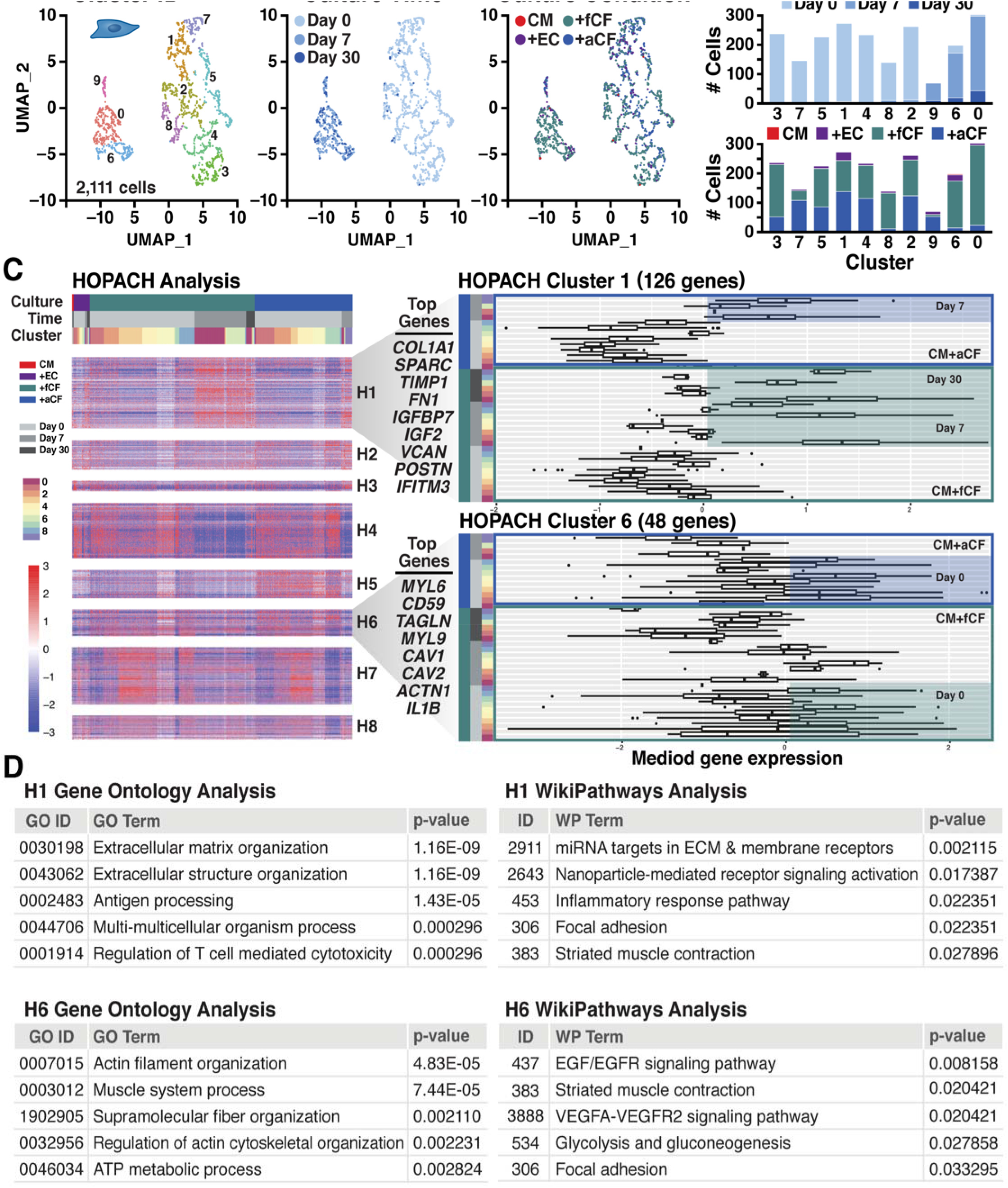
Cardiac fibroblast subset analysis of single-cell RNA-sequencing data. (A) UMAP clustering of classified CFs colored by cluster identity, culture time point, and tissue condition. (B) Cell counts for each UMAP cluster organized by time point and by tissue condition. (C) Heatmap of HOPACH clustering of CFs and plot of HOPACH clusters 1 and 6 (H1 and H6, respectively). (D) Gene ontology and WikiPathways results for H1 and H6.

Cluster analysis revealed 10 individual subsets of fibroblasts, with CF expression patterns after 7 and 30 days of microtissue culture (UMAP clusters 0, 6, and 9) clearly distinct from the starting CF populations (Figure 9A). HOPACH analysis identified 8 clusters of genes (H1_CF_-H8_CF_) that drove phenotypic differences in CFs as a result of time in culture (day 0, day 7, day 30) and developmental stage (fCF vs. aCF). H1_CF_ contained 126 genes that were highly expressed in fCFs and aCFs after 3D heterotypic culture (days 7 and 30), while H6_CF_ contained 48 genes that were upregulated in input (day 0) CFs (Figure 9C). The H1_CF_ genes *(COL1A1, SPARC, TIMP1, FN1)* are related to extracellular matrix organization and antigen processing, while H6_CF_ genes *(MYL6, TAGLN, MYL9)* are related to cytoskeletal organization and muscle contraction (Figure 9D). Differences between CFs at distinct developmental stages were reflected in H5CF, which contained 51 genes (including *TPM1*, *MYL12A*, and *IL1B*) that were expressed at higher levels in day 0 adult CFs than day 0 fetal CFs, with GO terms related to muscle contraction and actin organization (Supplementary Figure 4). The remaining HOPACH clusters listed differentially expressed genes that were not specific to CFs (Supplementary Figure 4, Supplementary Table 3). In summary, this data reveals that 3D cardiomyocyte co-culture induces a less contractile fibroblast phenotype and promotes the fibroblasts’ ability to remodel their extracellular environment.

## Discussion

This study interrogated the heterotypic influences of paired cardiac populations on cellular phenotype and function within 3D engineered cardiac tissues. Due to the single cell-based nature of the analyses, we were able to dissect bi-directional phenotypic changes in each specific cellular sub-type and evaluate them in the context of tissue level functional performance. While it is well established that non-myocytes are required to promote stable tissue formation^21,26–29^, previous studies have described variable effects of different stromal or non-myocyte populations on cardiac function with limited details reported on phenotypic shifts in the co-cultured cells^21,30,31^. Therefore, we aimed to determine how age-specific CFs and ECs differentially affect cardiac tissue organization and calcium handling dynamics, resulting in transcriptional changes in the different heterotypic populations of cells.

hiPSC-CMs lack a fully mature phenotype, and transcriptional progression throughout differentiation renders them most comparable to early-/mid-gestation fetal CMs^32–34^. Several key environmental parameters have been implicated in promoting the developmental phenotype of hiPSC-CMs, such as three-dimensional culture^35–38^, biochemical cues^39–41^, electromechanical stimulation^37,38,42–45^, and extended culture duration^46–48^. Consistent with previous studies, the cardiac microtissues examined here exhibited overall improvements in calcium handling properties (amplitude, maximum upstroke and downstroke velocities) with prolonged (30 day) culture in the engineered 3D platform. However, despite more mature calcium transient profiles, the day 30 microtissues could not respond to the higher stimulation frequencies, potentially indicating that since the microtissues were able to take up more calcium (increased amplitude) they might not have been able to release it in time, and therefore were still in a refractory phase when the faster stimulation pulses were fired. Overall, the functional changes at day 30 occurred in all microtissues regardless of the heterotypic pairing, highlighting culture duration aids in modest CM maturation.

In contrast, short-term co-culture (first 7 days) with CFs promoted more rapid phenotypic maturation of CMs, in terms of both microtissue calcium handling dynamics and CM gene expression. Our finding that co-culture with ECs did not improve cardiac microtissue function is consistent with previous reports showing that while CMs may be transcriptionally or structurally impacted by EC co-culture, functional parameters are not improved^49,50^. However, a recent study demonstrated that ECs can improve stem cell-derived cardiac microtissue function when they are incorporated with a stromal population (i.e. CM+CF+EC microtissues), indicating a potential additive effect of heterotypic tri-culture on tissue function^31^.

After 7 days of culture, cytoskeletal *(RGS5, PDLIM3)* as well as ECM-associated *(COL3A1, IGF2, SPARC, TIMP1)* genes were more highly expressed in CMs cultured with CFs than in CMs cultured alone or with ECs. Notably, both *RGS5* and *PDLIM3* are implicated in cardiac homeostasis and calcium handling. *RGS5,* a regulator of G protein signaling that is expressed in multiple cardiac cell types throughout development, protects CMs from apoptosis, inflammation, and fibrotic remodeling^51,52^. Moreover, a mouse model of *RGS5* deficiency exhibits prolonged cardiac repolarization and increased action potential duration^53^, indicating a role for *RGS5* in cardiac electrophysiology. *PDLIM3* (PDZ and LIM Domain 3), also known as actin-associated LIM protein or ALP, works in concert with muscle LIM protein (MLP), by co-localizing to the intercalated disks and interacting with alpha actinin^54^, contributing to cardiac muscle organization and maintenance. Absence or dysfunction of ALP and MLP are associated with heart disease^55,56^, and MLP-deficient hPSC-CMs display impaired calcium handling and progressively mimic hypertrophic cardiomyopathy^57^. Therefore, the roles of *RGS5* and *PDLIM3* in cytoskeletal structure and cardiac function are consistent with our finding that increased expression of those genes in CMs paired with CFs correlated with improved calcium handling properties.

Throughout heart development and in response to injury, CFs are considered the primary cell type responsible for ECM synthesis and remodeling^15,58^. Consistent with this paradigm, we observed increased expression of several genes related to ECM-production (*COL1A1*, *FN1*, *VCAN*, *POSTN*) and ECM-remodeling (*TIMP1)* in CFs from microtissues. Notably, however, CMs co-cultured with CFs also exhibited increased expression of ECM-related genes (e.g. *COL3A1* and *TIMP1),* suggesting that CMs may also actively modulate their extracellular microenvironment in the presence of CFs.

Endothelial cells were also impacted by 3D heterotypic co-culture, though the effects were harder to dissect due to low numbers of cells in the day 7 and day 30 sequencing analysis as well as the identified cells that came from CF microtissues, which could be due to the impure nature of primary CF isolations or a result of the bioinformatic cell “sorting” pipeline. Cell classification was based on expression of cell-specific genes, but the lack of definitive non-myocytes markers makes the bioinformatic pipeline vulnerable to the same potential flaws of physical cell sorting, where there is not enough stark contrast between the non-myocytes to accurately distinguish ECs and CFs. HOPACH analysis of the EC subset identified clusters of genes that aligned with tissue culture time points: H1EC was driven by input (day 0) ECs (98% from +EC tissues) and corresponded to metabolic-related GO and Wikipathways terms, while H3_EC_ and H4_EC_ were driven by day 7 and day 30 ECs (~22% from +EC tissues, 78% from +CF tissues), and were associated with endothelial development and cardiac calcium regulation (H3_EC_) and immune response (H4EC) terms. It is difficult to interpret these data due to the relatively large presence of ECs in the +fCF and +aCF microtissues, but the genes that informed the cardiac calcium regulation WikiPathways terms in H3_EC_ included Connexin 37 *(GJA4)* and Calmodulin 1 (*CALM1*), both of which are known regulators of ion transport. Gap junction GJA4 is most commonly found between endothelial cells, and both CALM1 and GJA4 interact with endothelial nitric oxide synthase (eNOS), which synthesizes the nitric oxide that ECs secrete for CM homeostasis and contractility^9,59,60^. Therefore, the presence of these genes may indicate that 3D heterotypic culture with CMs promotes paracrine signaling in ECs.

In the context of this study, there were few discernable differences in the performance or phenotype of microtissues comprised of CMs with either fetal or adult CFs, indicating that the temporal stage of a specific non-myocyte population does not impact phenotypic and functional differences as much as the type of non-myocyte. We observed a set of genes expressed in input adult CFs that were not expressed in input fetal CFs, but after mixing with CMs in microtissue culture, these differences largely disappeared and the CFs were more phenotypically similar to one another. CM+fCF microtissues maintained a core of fibroblasts throughout the 30 days of culture and were able to maintain 2Hz electrical pacing at day 30, whereas the non-myocyte cores within CM+aCF tissues became more diffuse by day 30 and the microtissues were not able to sustain pacing >1Hz. This may indicate that the persistent presence of a non-myocyte core is key to improved pacing response. Furthermore, in day 7 microtissues, the expression of cTnI was elevated at the borders of the fCF and aCF cores, indicating that CMs juxtaposed with CFs may mature faster. The expression of increased by day 30 in all conditions, consistent with the paradigm that culture duration increases maturation, and with the finding that day 30 tissues had improved calcium handling compared to their day 7 counterparts. The ability of CFs to act as electrical insulators or provide impulse propagation and excitability to their neighbors remains debated in the field, though a recent study demonstrated that Connexin 43 gap junctions between CMs and CFs are necessary for microtissue function and maturation^31^. Advanced spatial transcriptomic and 3D imaging technologies^61^ will provide further insight into the influence of multicellular spatial organization on beat rate, pacing responses, and intratissue heterogeneity.

Previous characterization of cardiac microtissue heterogeneity using light sheet microscopy determined that the initial seeding ratio of 3:1 CMs:CFs is maintained after 7 days of microtissue culture^61^, which suggests that the lower numbers of ECs and CFs identified at day 7 in the single-cell RNA-seq analysis may be due to a technical artifact. Low recovery of non-myocytes could result from dissociation removing cells in radial manner, such that the central core of CFs are last and hardest to dissociate. However, the number of identified non-myocytes was even further diminished in day 30 heterotypic cultures, indicating that this experimental system favors CMs. This further decline may be related to the use of hiPSC-CM maintenance medium, and therefore media optimization may be required to support non-myocyte survival and/or function long term^31^. Morphologically, we observed that the frequency of non-myocyte cores decreased by day 30, and functionally, day 30 heterotypic microtissues displayed less mature calcium handling properties than the homotypic microtissues. Therefore, it remains unclear whether the positive contribution of CFs was a transient phenomenon, or if the progressive loss of non-myocytes with long-term culture adversely affected microtissue function.

Increasing complexity of microtissues by incorporation of multiple cardiac cell types, modified tissue geometries, and the addition of biophysical or electromechanical cues will advance understanding of the complex multisystem crosstalk in these systems. This study demonstrates how single-cell phenotypic analysis can yield a deeper understanding of how heterotypic interactions affect individual cell phenotypes and their collective contribution to tissue-level structural and functional properties. The ability to merge single-cell metrics with multicellular tissue behaviors is paramount to developing advanced systems capable of modeling human development and disease *ex vivo.*

## Methods

### Pluripotent stem cell culture

WTC11 human induced pluripotent stem cells (hiPSCs) modified with a genetically-encoded calcium indicator GCaMP6f^62,63^ (generously donated by Dr. Bruce Conklin) were cultured in mTeSR medium (Stem Cell Technologies, Vancouver, CA) on plates coated with 80μg/mL Matrigel (Corning, Corning, NY). Cells were seeded at 1×10^4^ cells/cm^2^ in mTeSR medium, supplemented with 10μM Rock inhibitor (Y27632, SelleckChem, Houston, TX) for the first 24h, and passaged using Accutase (Innovative Cell Technologies, San Diego, CA) every 3 days at ~70% confluence.

### Cardiomyocyte differentiation

Cardiomyocytes (CM) were differentiated from WTC11 GCaMP6f hiPSCs following a chemically-defined, serum-free protocol^64,65^. Cells were seeded onto Matrigel-coated 12-well tissue culture plates at 3×10^4^ cells/cm^2^ in mTeSR medium with 10μM Rock inhibitor. Once the cells reached 100% confluence, 12μM CHIR99021 (SelleckChem) was added into RPMI 1640 medium (Thermo Fisher, Waltham, MA) supplemented with B27 minus insulin (denoted as RPMI/B27-; Life Technologies, Grand Island, NY) (differentiation day 0). CHIR was removed exactly 24h later by replacing medium with fresh RPMI/B27-. On day 3, 5μM IWP2 (Tocris, Bristol, UK) in RPMI/B27-was added and then removed 48h later by refreshing the medium. The cells were fed with fresh RPMI/B27-on day 5. On day 7, medium was switched to RPMI 1640 medium supplemented with B27 with insulin (RPMI/B27+; Life Technologies) and fed every 3 days thereafter. On day 15, cells were re-plated onto Matrigel-coated dishes at 1×10^5^ cells/cm^2^ in RPMI/B27+ with 10μM Rock inhibitor. Purification of cardiomyocytes occurred by feeding cultures with Lactate purification medium^66^ (no-glucose Dulbecco’s Modified Eagle Medium (Thermo Fisher) with 1X Non Essential Amino Acids (NEAA; Corning), 1X Glutamax (*L*-glut; Life Technologies), and 4mM Lactate) on days 20 and 22. On day 24, medium was exchanged to RPMI/B27+ and fed every three days thereafter until harvest (day 35 ± 7).

### Endothelial cell differentiation

Differentiation of endothelial cells from hiPSCs was achieved by modifying a previously described protocol^67^. Briefly, on day 0, WTC11 hiPSCs were seeded onto Matrigel-coated dishes in E8 medium (Thermo Fisher) supplemented with 5ng/mL BMP4 (Stem Cell Technologies), 25ng/mL Activin A (Stem Cell Technologies), and 1μM CHIR99021, with daily media changes. On day 2, cells were cultured in E6 medium (Thermo Fisher) supplemented with 100ng/mL bFGF (Stem Cell Technologies), 50ng/mL VEGF-A (Stem Cell Technologies), 50ng/mL BMP4, and 5μM SB431542 (Stem Cell Technologies), with daily media changes. On day 4, cells were split onto a fibronectin (Sigma-Aldrich, St. Louis, MO) coated plate and maintained in EGM medium (Lonza, Basel, SUI). The efficiency of the differentiation was assessed on day 8 with flow cytometry using CD31 and CD144 antibodies, and was typically >95% CD31+/CD144+.

### Fibroblast cell culture

Human fetal (18wk gestation, male, lot #2584) and adult (50 y.o., male, lot #3067) cardiac fibroblasts (Cell Applications, San Diego, CA) were cultured according to manufacturer recommendations. Briefly, cells were seeded and passaged at a density of 1×10^4^ cells/cm^2^, and fed with Cardiac Fibroblast Medium (Cell Applications) every 2-3 days, for up to 10 passages.

### Cardiac microtissue formation

Lactate-purified cardiomyocytes (day 35 ± 7), fetal cardiac fibroblasts, adult cardiac fibroblasts, and endothelial cells were dissociated with 0.25% Trypsin for 5-10 min and then mixed together at a 3:1 ratio of CM to nonmyocytes. Cell mixtures in RPMI/B27+ with 10μM Rock inhibitor were seeded into 400μm inverted pyramidal agarose microwells at a density of 2000 cells/well and allowed to self-assemble overnight^26^. After 18-24h, the self-assembled cardiac microtissues were removed from the wells and maintained in rotary suspension culture at a density of 2000 tissues per 10cm Petri dish for 7 or 30 days.

### Histology and immunofluorescence staining

Samples were fixed in 10% Neutral Buffered Formalin (VWR, Radnor, PA) for 1h at room temperature and embedded in HistoGel Specimen Processing Gel (Thermo Fisher) prior to paraffin processing. Five micron sections were cut and adhered to positively charged glass slides. Slides were deparaffinized with xylene and re-hydrated through a series of decreasing ethanol concentrations (100%, 100%, 95%, 80%, 70%). For immunofluorescence staining, epitope retrieval was performed by submersing slides in Citrate Buffer pH 6.0 (Vector Laboratories, Burlingame, CA) in a 95°C water bath for 35min. Slides were cooled at RT for 20 min and washed with PBS. Samples were permeabilized in 0.2% Triton X-100 (Sigma-Aldrich) for 5min, blocked in 1.5% normal donkey serum (Jackson Immunoresearch, West Grove, PA) for 1h, and probed with primary and secondary antibodies against cardiac troponin T, slow+fast troponin I, and cardiac troponin I, and counterstained with Hoechst and WGA (antibody information in Supplementary Table 4). Coverslips were mounted with anti-fade mounting medium (ProlongGold, Life Technologies) and samples were imaged on a Zeiss Axio Observer Z1 inverted microscope equipped with a Hamamatsu ORCA-Flash 4.0 camera.

### Calcium imaging analysis

Calcium transients of cardiac microtissues were visible due to the genetically-encoded calcium indicator GCaMP6f in the hiPSC-CMs. Microtissues at day 7 or day 30 were incubated in Tyrode’s solution (137 mM NaCl, 2.7 mM KCl, 1 mM MgCl2, 0.2 mM Na2HPO4, 12 mM NaHCO3, 5.5 mM D-glucose, 1.8mM CaCl2) in a 35mm Petri dish for 30min at 37°C prior to imaging. Samples were mounted on a Zeiss Axio Observer Z1 inverted microscope with a Hamamatsu ORCA-Flash 4.0 camera. Electrodes were placed in the Petri dish to apply electrical field stimulation at 1Hz as well as a stimulation regimen of increasing frequencies (0.5Hz, 1Hz, 2Hz, 4Hz) (MyoPacer, IonOptix). Calcium transient videos were acquired with Zen Professional software (v.2.0.0.0) at 10ms exposure and 100 frames per second. Circular regions of interest (ROI; 75-pixel diameter) were selected at the center of each microtissue and the mean fluorescent intensity values were plotted against time. Metrics of calcium transient kinetics, such as amplitude, stroke velocities, and beat rate, were analyzed using a custom python script^68^. Source code is available at https://github.com/david-a-joy/multilineage-organoid.

### Statistics

The mean +/− standard deviation was calculated from at least 9 biological replicates for all data unless otherwise noted. When comparing three or more groups, one-way analysis of variance (ANOVA) followed by Tukey’s post hoc analysis was performed. For all comparisons, statistical significance was determined at p<0.05. All statistical analysis was performed using GraphPad Prism 7.0 software.

### Single-cell RNA-sequencing sample preparation and sequencing

At the time of cardiac tissue formation (day 0), a subset of dissociated heterotypic cell mixtures was retained for sequencing. After 7 or 30 days of microtissues culture, cells were dissociated with 0.25% Trypsin for 45 minutes and ~8000 cells/group) were prepared for analysis through droplet encapsulation by the Chromium Controller and library preparation with the Chromium Single Cell 3’ v2 Library and Gel Bead Kit (10x Genomics, San Francisco, CA). cDNA was sheared using a Covaris S2 sonicator and 12 PCR cycles were run during cDNA amplification. Libraries were sequenced on a HiSeq 4000 (Illumina, San Diego, CA). Sequences were demultiplexed and aligned to human reference genome grch38 using the default settings of 10x Genomics *CellRanger* v2.0.2. After *CellRanger* filtering, there were ~700 million valid reads, 88.6% of which were mapped to a unique UMI. Raw data used for single-cell RNA-sequencing analysis is uploaded to GEO under the accession number (upload in progress).

### Single-cell RNA-sequencing zinbwave reduction

The raw counts for genes from all cells across culture conditions and time points were loaded with Seurat v2^69,70^. The cells were filtered using the number of detected genes between the bottom 1% and top 99% across all cells, and percent mitochondrial genes within the top 99% across all cells. Genes were filtered out of further analyses if they did not have at least 5 counts in at least 5 cells. The zinbwave function (with parameter K=2) in the R bioconductor package zinbwave was used to generate a 2-dimensional (2D) representation of the gene expression profiles for each of the cells after adjusting for the number of detected genes per cell^71,72^. The zinbwave function (with parameter K=0 and adjusting for the number of genes detected per cell) was used to generate weights per gene-cell combination that were used in the gene expression association analyses with edgeR^73,74^. The 2D zinbwave reduction of all of the cells was visualized as a umap^75^. The resulting zinbwave reduction was loaded as a Seurat v2 object using SetDimReduction function. Clustering of cells was performed using the FindClusters function in Seurat v2 using parameter resolution=0.6. Cells in clusters expressing *TNNT2*, *TNNI1*, and *ACTC1* were chosen for the CM-specific analyses, cells in clusters expressing *POSTN, THY1,* and *FN1* were chosen for the CF-specific analyses, and cells in clusters expressing *PECAM1*, *KDR*, and *FLT1* were chosen for the EC specific-analyses. The raw counts from the resulting cells of an identified cell type (CM, CF, or EC) were used to generate the 2D representation of the gene expression profiles as before, using the zinbwave function. The resulting zinbwave reductions were loaded as a Seurat v3 object using the as.Seurat function. Clustering of each cell type was performed using the FindClusters function in Seurat v3 using parameter resolution=0.4. Genes whose expression was associated with culture condition and/or time were determined using edgeR. Weights associated with gene-cell combinations that were estimated from running the zinbwave functions were used as weights in the edgeR-specific DGE object. The design matrix used for the all-cell analyses (pre-cell type subsetting) for each gene’s expression was: Y ~ nGene + Clusters + Culture + Time + Clusters:Culture + Culture:Time + Clusters:Time. The design matrix for the cell-type specific analysis was: Y ~ nGene + Clusters + Time + Clusters:Time. Associated genes were determined by the significance (FDR < 0.05) of the composite null hypothesis involving terms containing culture conditions and/or time. Normalized expression for all genes meeting the significance threshold over all cells was obtained from the computeDevianceResiduals function in zinbwave.

### Single-cell RNA-sequencing HOPACH analysis

The normalized gene expression data were used to cluster genes across all of the cells using Hierarchical Ordered Partitioning and Collapsing Hybrid (HOPACH) analysis^76^. HOPACH determined the optimal number of gene clusters that best described the given data as well as the identity of the genes belonging to each HOPACH cluster. The normalized expression data was visualized as a heatmap using the pheatmap package^77^. Gene clusters were also visualized as box plots of expression of medoid (representative) genes from each HOPACH cluster, based on groups of cells belonging to a particular combination of culture condition, time-point and cell-cluster membership. GO analysis and WikiPathways analysis were performed on the HOPACH cluster gene lists using the Cluster Profiler R package^78^.

### In situ hybridization

RNAScope for *PDLIM3, RGS5, COL3A1,* and *IGF2* (probe information in Supplementary Table 4) was performed on sections of formalin-fixed paraffin-embedded samples (see “histology and immunofluorescence staining”) using the RNAscope Multiplex Fluorescent Reagent Kit v2 (Advanced Cell Diagnostics, Newark, CA) and following the protocol outlined in User Manual 323100-USM. Sections were imaged on a Zeiss Axio Observer Z1 inverted microscope equipped with a Hamamatsu ORCA-Flash 4.0 camera.

## Acknowledgements

The authors acknowledge funding support from the California Institute for Regenerative Medicine (LA1-08015), the NSF Engineering Research Center for Cell Manufacturing Technologies (CMaT; NSF EEC-1648035), and the Gladstone BioFulcrum Heart Failure Research Program. T.A.H. was supported by an American Heart Association Postdoctoral Fellowship (15POST22750003). O.B.M. is a National Science Foundation Graduate Research Fellow (1650113). The authors would like to thank the Gladstone Bioinformatics Core, Gladstone Genomics Core (Dr. Natasha Carli and Jim McGuire), and the Gladstone Stem Cell Core (Dr. Po-Lin So; Roddenberry Stem Cell Foundation) for assisting with experimental analysis. The authors would also like to thank Dr. Nathaniel Huebsch and Dr. Bruce Conklin for providing the WTC11-GCaMP hiPSC line, Dr. Sanjeev Ranade for assisting with the hiPSC-EC differentiations, and Dr. Michael Lai for relevant critical discussion. Many thanks to the Gladstone art department (Giovanni Maki and Sarah Gardner) for assistance with graphics and Dr. Kathryn Claiborn for reviewing the manuscript and scientific editing.

## Author Contributions

T.A.H., O.B.M, and T.C.M. designed the experiments; T.A.H., O.B.M, and J.E.S. performed the experiments; D.A.J. developed the python script for calcium imaging analysis; R.T. performed single-cell RNA-seq zinbwave reduction and HOPACH analysis; T.A.H., O.B.M., D.A.J., and R.T. performed single-cell RNA-seq analysis. T.A.H., O.B.M., and T.C.M. wrote the manuscript.

