## Supplementary figures and tables for "Bi-directional Impacts of Heterotypic Interactions in Engineered 3D Human Cardiac Microtissues Revealed by Single-Cell RNA-Sequencing and Functional Analysis"

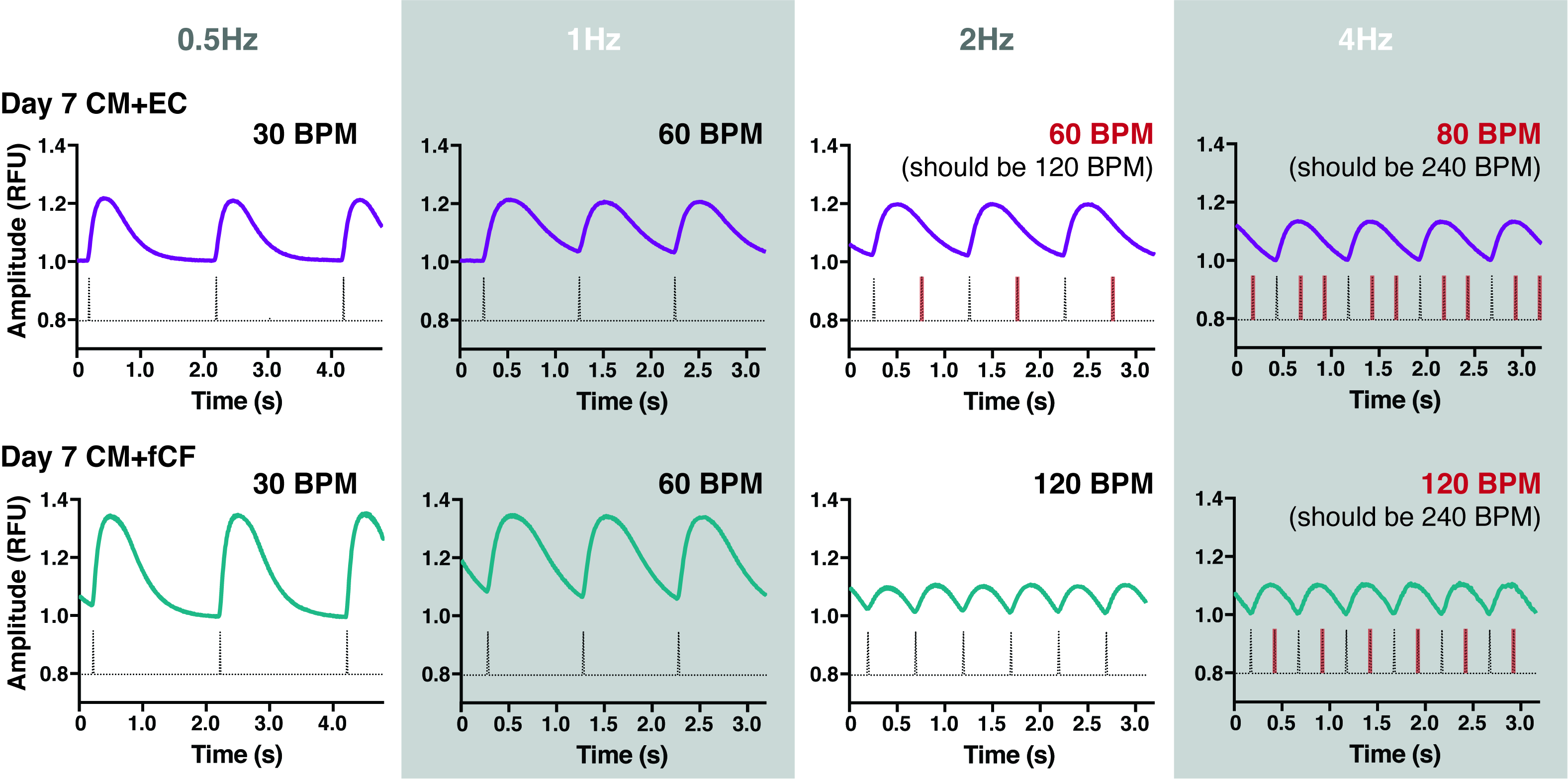

**Supplementary Figure 1.** Calcium transient traces aligned with pulse train of stimulation frequencies for representative day 7 CM+EC (top) and CM+fCF (bottom) microtissues. Red lines on pulse train indicate a missed beat. CM+EC microtissue did not responded appropriately at 2Hz and 4Hz stimulation while CM+fCF microtissue did not responded to 4Hz stimulation.

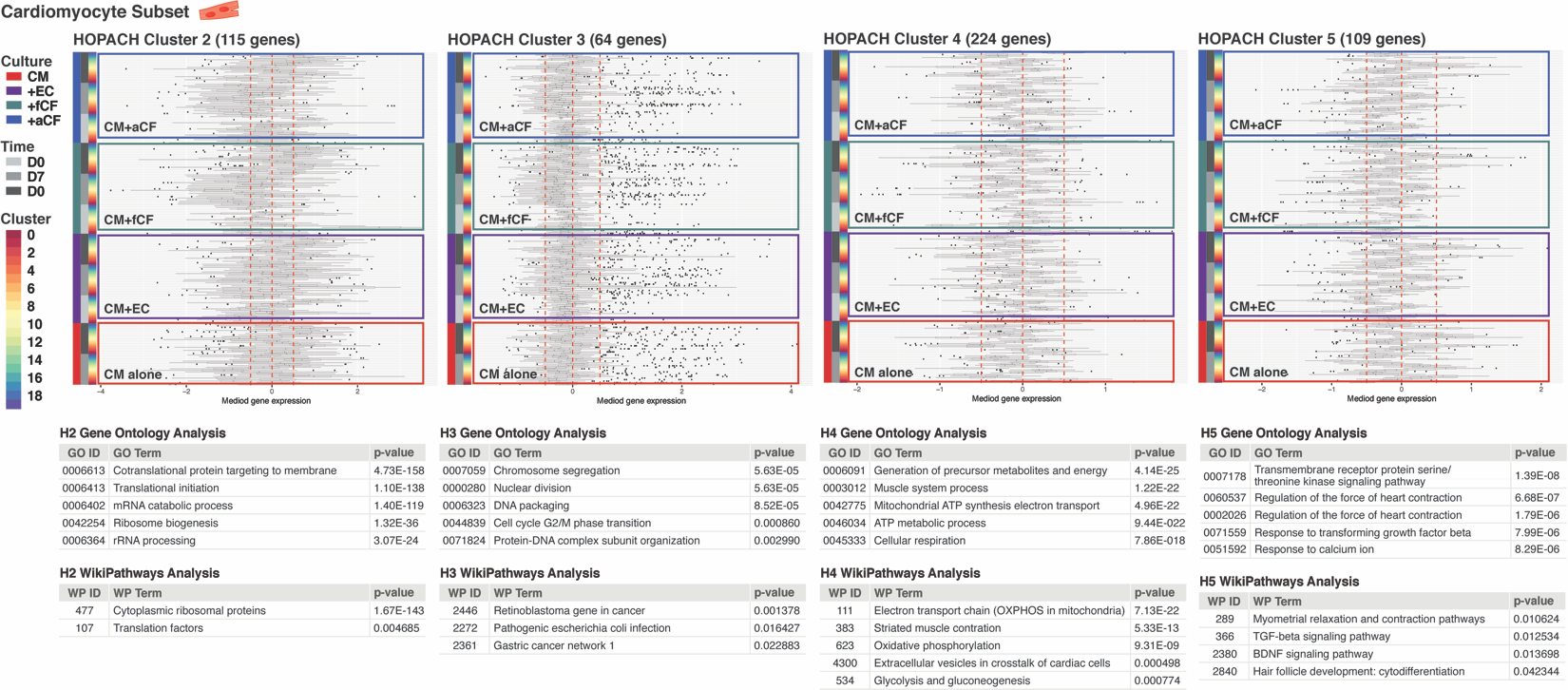

**Supplementary Figure 2.** Remaining HOPACH clusters (H2-H5) for CM subset analysis with corresponding gene ontology and WikiPathways results.

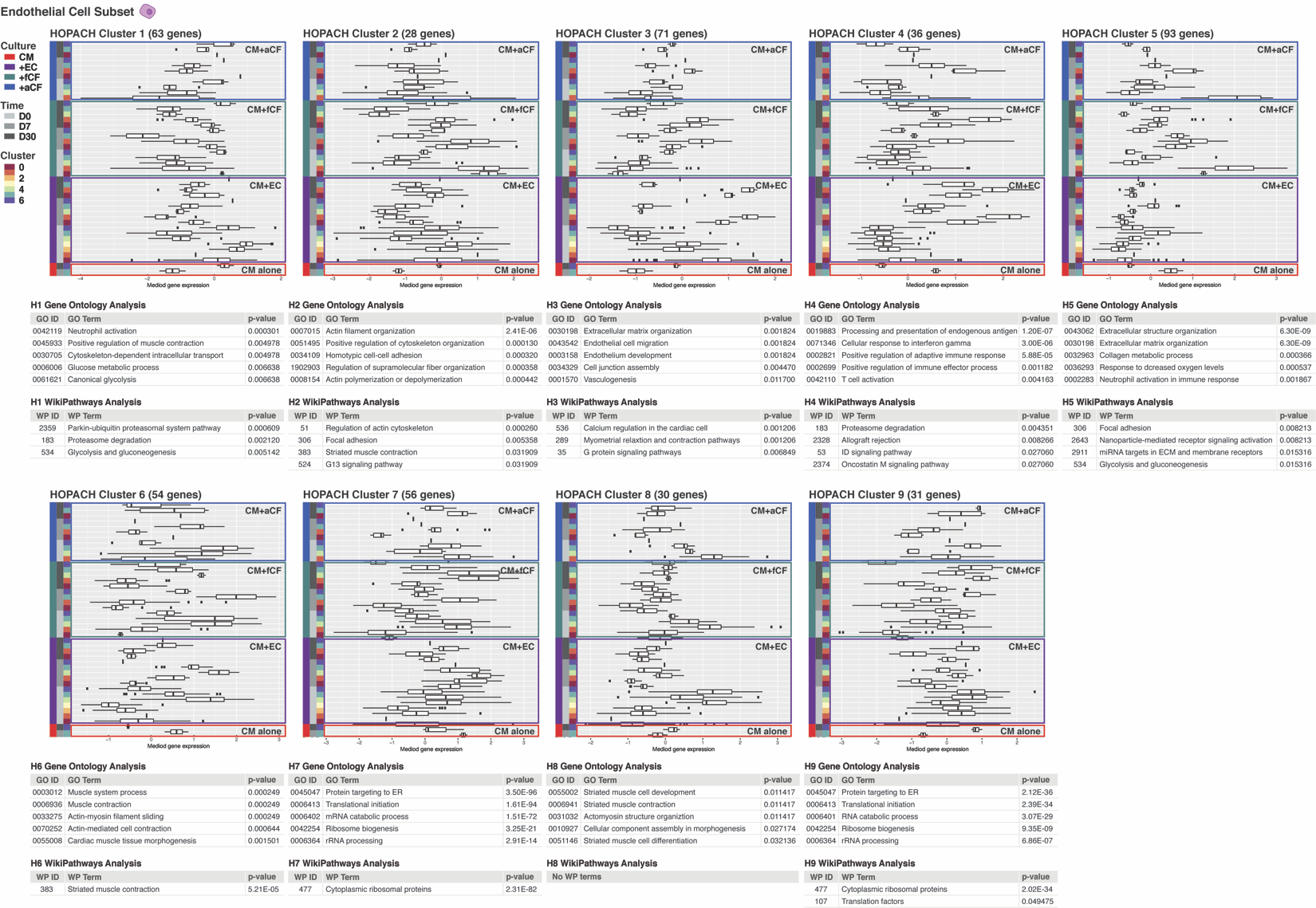

**Supplementary Figure 3.** All HOPACH clusters (H1-H9) for EC subset analysis with corresponding gene ontology and WikiPathways results.

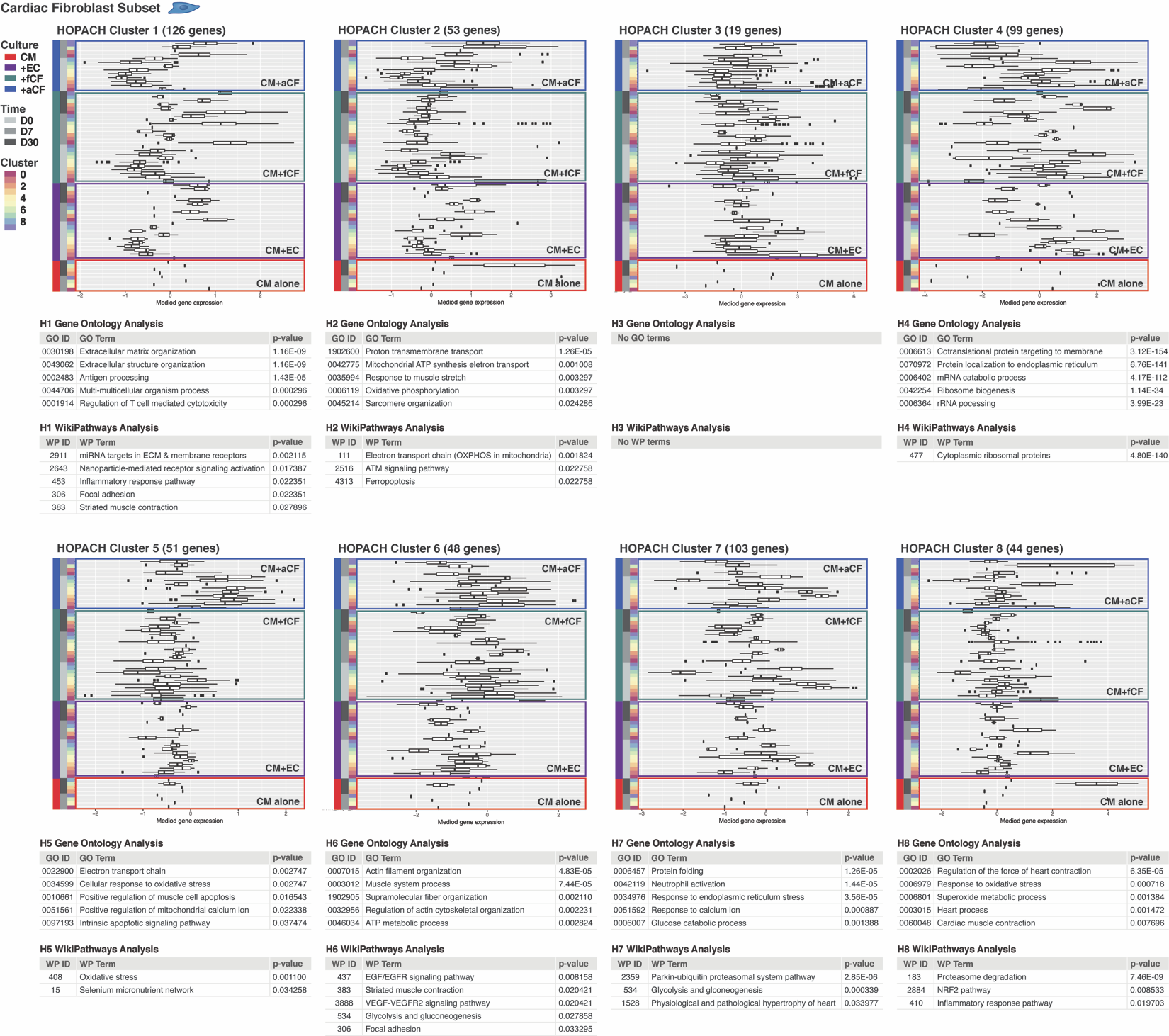

**Supplementary Figure 4.** All HOPACH clusters (H1-H8) for CF subset analysis with corresponding gene ontology and WikiPathways results.

**Supplementary Table 4.** Antibody and in situ hybridization probe source information.

| **Immunocytochemistry** | | | | | | | |
| --- | --- | --- | --- | --- | --- | --- | --- |
| **Antibody** | | | **Company** | | **Catalog #** | | **Dilution** |
| Cardiac Troponin T | | | Abcam | | ab45932 | | 1:400 |
| Wheat Germ Agglutinin | | | Thermo Fisher | | W11263 | | 1:500 |
| Slow+Fast Troponin I | | | Abcam | | ab47003 | | 1:100 |
| Cardiac Troponin I | | | Abcam | | ab110132 | | 1:100 |
| Alexa Fluor 555 | | | Thermo Fisher | | A-31572 | | 1:400 |
| Alexa Fluor 488 | | | Thermo Fisher | | A-21202 | | 1:400 |
| Hoechst | | | Thermo Fisher | | 62249 | | 1:10000 |
| **RNAscope in situ hybridization (Advanced Cell Diagnostics)** | | | | | | | |
| **Gene** | **Species** | **Target Region** | | **Amplification Channel** | | **Catalog #** | |
| *PDLIM3* | Hs | 254-1190 of  NM_014476.5 | | C3 | | 533411 | |
| *RGS5* | Hs | 339-1572 of  NM_003617.3 | | C3 | | 533421 | |
| *COL3A1* | Hs | 3550-5057 of  NM_000090.3 | | C1 | | 549431 | |
| *IGF2* | Hs | 339-1572 of  NM_003617.3 | | C2 | | 594361 | |
